# Targeted medial prefrontal cortex stimulation prevents incubation of cocaine craving and restores functional connectivity

**DOI:** 10.64898/2026.04.21.719530

**Authors:** Hanbing Lu, Samantha Hoffman, Ying Duan, Zilu Ma, Hieu Nguyen, Aidan Carney, Taylor Scott, Olivea Varlas, Md Mohaiminul Haque, Elliot A. Stein, Zheng-Xiong Xi, Yavin Shaham, Yihong Yang

## Abstract

**Background:** Relapse remains a central obstacle in the treatment of cocaine use disorder (CUD), for which no medications have received approval from the U.S. Food and Drug Administration. Transcranial magnetic stimulation (TMS) has shown promise as a potential therapeutic intervention. However, current clinical trials often rely on a “trial-and-error” approach in target selection and experimental design. We previously developed a novel TMS platform and high-density theta burst stimulation (hdTBS) technology, enabling precise, focal stimulation of the rat medial prefrontal cortex (mPFC), including the prelimbic and anterior cingulate cortices.

**Methods:** We applied hdTBS intervention to a well-established rat model of cocaine relapse and craving after cessation of extended access intravenous drug self-administration and assessed brain response using resting-state functional magnetic resonance imaging (fMRI).

**Results:** As expected, we observed robust time-dependent increases in cocaine seeking (incubation of cocaine craving) in control rats receiving sham stimulation over 3 weeks of abstinence accompanied by a reduction in prefrontal functional connectivity. In contrast, daily sessions of hdTBS for 7 days delivered on abstinence days 14-20 prevented the emergence of the incubation effect and restored prefrontal network functional connectivity.

**Conclusions:** This study provides strong preclinical evidence demonstrating that precise circuit modulation of medial prefrontal subregions causally reverses both behavioral and network-level adaptations associated with relapse vulnerability. Given the clinical accessibility and established safety profile of TMS, this work provides a mechanistically grounded framework for target selection and supports the translation of focal TMS of the mPFC for relapse prevention in CUD patient.

**One Sentence Summary:** Focal transcranial magnetic stimulation (TMS) of the mPFC using the hdTBS procedure prevents incubation of cocaine craving and restores functional connectivity.

## INTRODUCTION

Cocaine use remains a major public health concern, impacting an estimated 22 million people globally (1). In the United States, deaths from cocaine-related overdoses have increased more than fourfold over the past decade (2). There is currently no medication approved by the U.S. Food and Drug Administration for the treatment of cocaine use disorder (CUD). Recently, brain stimulation has been proposed as a novel treatment strategy aiming to correct circuit function changes hypothesized to be implicated in addiction (3). However, clinical trials involving techniques such as deep brain stimulation (4), transcranial direct current stimulation and transcranial magnetic stimulation (TMS) have followed a trial-and-error approach, yielding mixed results (5–7). Preclinical brain stimulation studies are essential for guiding and refining these interventions to enhance their therapeutic potential.

Relapse to drug use, even after prolonged abstinence, remains a major challenge in CUD treatment (8). In humans, drug craving and relapse are often triggered by re-exposure to environmental cues and contexts previously associated with drug use (9). This phenomenon has been modeled in rats with a history of cocaine self-administration, where cue-induced drug seeking intensifies, or “incubates,” during a period of forced abstinence (10–12). Several human studies specifically designed to test incubation of cue-induced drug craving reported findings consistent with preclinical studies (13–15). Notably, the abstinence phase in this model provides a window for repeated TMS sessions, which are necessary to achieve therapeutic effects. This makes the incubation model especially valuable for evaluating TMS as a therapeutic intervention in preclinical models of relapse.

Applying TMS in rat disease models presents technical challenges related to coil focality. A small TMS coil is necessary to minimize off target effects (16), but is associated with low efficiency, coil overheating and excessive electromagnetic stress (17,18). Previous preclinical studies typically used anesthetized rodents with either oversized coils that stimulated large brain areas or miniaturized coils that were too weak to induce action potentials. To overcome these limitations, we designed and built a coil platform that enables highly focal (∼2mm), suprathreshold stimulation of specific brain regions with a focality comparable to that in humans (19). Additionally, we recently introduced a high-density theta burst stimulation (hdTBS) methodology which delivers 6 pulses per burst as opposed to only 3 pulses in conventional TBS (20), enhancing acute after-effects of TMS by 92% (21).

In this study, leveraging the focal TMS coil and hdTBS technology, we applied TMS to the dorsal medial prefrontal cortex (mPFC) of rats during abstinence from cocaine SA. We chose the mPFC because of its critical role in cocaine cue-induced reinstatement and incubation of cocaine seeking (22–26). We found that 7 daily hdTBS sessions during the abstinence phase reversed the incubation effect and restored incubation-associated decreases in functional connectivity, assessed by functional MRI. Our preclinical study demonstrates the therapeutic potential of TMS for cocaine addiction with coil focality and stimulation parameters closely mirroring those used in clinical applications. This methodological alignment significantly enhances the translational relevance of our findings.

## METHODS

### Subjects

Sprague-Dawley male rats (Charles River, n=33 total), aged 90-120 days (300-350 grams) were used. Rats were initially group-housed and then individually housed after surgery. They were maintained on a light-dark cycle with *ad libitum* access to food and water. Procedures were approved by NIDA Animal Care and Use Committee.

### Behavior experiments

#### Cocaine self-administration

As previously described (27), a customized catheter was implanted into the rat jugular vein under isoflurane anesthesia (3% induction; 1.5% to 2.5% maintenance). After 7-10 recovery days, rats were trained on a fixed-ratio 1 (FR1) 20-s timeout reinforcement schedule to self-administer cocaine (0.5 mg/kg per infusion over 3.5 s) for 3-h/day for 15 days in standard Med Associates SA chambers (Med Associates). Each session began with the illumination of a red house light followed by the insertion of an active lever. Responses on this lever resulted in one cocaine infusion that was paired with a 3.5-s compound tone+light cue. Inactive lever-presses were recorded but had no consequence. The number of infusions was limited to 50/session. During abstinence, the rats were kept in their home cage.

#### Relapse tests

Relapse to cocaine-seeking was assessed during early (Day 1) and late (Day 21) abstinence. The experimental conditions during testing were identical to the training phase, except that lever-presses did not result in cocaine delivery (extinction conditions).

### Behavioral data analysis

We analyzed behavioral data using Linear Mixed Effects (LME) modeling in R (28). For the relapse test data, TREATMENT (active vs. sham TMS) and TIME (abstinence day 1 vs. 21) were fixed effects and subject (rat) was a random effect, accounting for within subject correlation. Degrees of freedom were estimated using the Kenward–Roger method. Significant interactions were followed by Tukey post-hoc test. Data are presented as mean ± standard deviation (S.D.) and statistical significance was set at *p*<0.05.

### TMS administration

#### Headpost implantation for consistent coil placement

Supplemental Figures 1 and 2 illustrate the hdTBS paradigm and the stimulator system. As the homemade TMS coil features a focality of ∼2 mm, this high focality necessitates a method to consistently direct the hotspot of the TMS coil to the target region. As such, we implanted a headpost on the rat’s skull to serve as a reference to facilitate daily accurate coil positioning. Detailed procedures for headpost implantation were described previously(29). Briefly, a 3D-printed, T-shaped headpost was fixed onto the rat skull such that the end-pad of the headpost directed the coil hotspot to the targeted region (hindlimb motor cortex or the mPFC). A detachable coil guide of variable thickness can be used to further adjust coil positioning when necessary.

#### TMS Habituation

Following surgery, rats were allowed to recover for one week before undergoing 6-days of habituation to minimize stress during TMS administration. This involved holding the rats under the coil and playing pre-recorded hdTBS sounds for 5 min three times/day. After habituation, the rats showed reduced stress, characterized by absence of such stress signs as screaming, urination, defecation, and attempts to escape.

#### TMS Delivery

Following TMS habituation, the motor threshold was assessed (also in the hand-held awake state), to inform the intensity of TMS pulses. The details of the motor threshold measurement were previously described (21,29). Once determined, TMS power was set at 125% of motor threshold in the active TMS group (or maximal machine output if a rat’s motor threshold was too high). The rat’s head was placed immediately below the coil surface; in the sham group, TMS power was set at 75% motor threshold and the rat’s head was placed 4 cm beneath the TMS coil, resulting in an electric field inside the brain close to zero based on E-field mapping, while the acoustic sound level was similar. The rats’ ear canals were occluded manually during TMS administration to minimize both the acute and chronic effects of acoustic noise. Effectiveness of sound attenuation was confirmed by the absence of a startle response during TMS administration, which would otherwise occur frequently.

The hdTBS procedure delivers 1200 pulses within 200 s. To preclude coil overheating, we applied dry ice to cool the coil during stimulation and allowed 60 min of cooling between TMS deliveries to ensure the coil core was fully cooled. The rats received daily TMS sessions for 7 days (during abstinence days 14-20) and tested for relapse on day 21 without stimulation.

### fMRI experiments

The fMRI acquisitions were time-sensitive and required completion within 24 h following cocaine-seeking testing. However, our MRI scanner was temporarily unavailable due to equipment failure during a portion of these critical periods. Consequently, of the 33 rats that completed the behavioral protocol, 19 underwent two MRI scans (n = 10, active hdTBS; n = 9, sham hdTBS).

#### Animal preparation

A combination of isoflurane and dexmedetomidine was used to anesthetize rats during fMRI experiments. Detailed procedures were previously described(30,31). Briefly, rats were initially anesthetized using 2.5% isoflurane in oxygen enriched air (70% N_2_ + 30% O_2_), followed by a bolus injection of dexmedetomidine (0.015 mg/kg, intraperitoneal). Rats were next transferred and head-fixed onto a customized MRI-compatible holder and isoflurane concentration gradually lowered and maintained at 0.5%-0.75% through a nose cone. Isoflurane and expired CO_2_ were actively scavenged through a vacuum line. The flow rates of the inlet and outlet gases in the nose cone were balanced. Continuous subcutaneous infusion of dexmedetomidine (0.015 mg/kg/h) was delivered via an infusion pump (PHD 2000, Harvard Apparatus). Blood oxygenation and respiration rate were monitored (Small animal Monitoring and Gating System, SA Instruments) to ensure stable physiology during fMRI recording, which occurred at least 60 min after anesthesia introduction (30). Body temperature was maintained by a temperature-controlled water-heating pad during imaging. The oxygen concentration in gas mixture was adjusted to ensure SPO_2_ of at least 95%.

#### MRI scan

MRI data were acquired with a Bruker Biospin 9.4T scanner running the Paravision 7.0 software platform (Bruker Medizintechnik). A volume quadrature transmit coil (model: MT0381) was used for radio frequency excitation, and a circular surface coil (model: MT0105-20) for MR signal reception. The decussation of the anterior commissure (∼−0.36 mm from bregma) served as the fiducial landmark to standardize slice localization both within and across rats (32).

#### Structural MRI scan

High-resolution structural images were collected using a Rapid Acquisition with Relaxation Enhancement (RARE) sequence. Scan parameters: repetition time (TR)=3100 ms, echo time (TE)=36 ms, field of view (FOV)=30×30 cm^2^, in-plane matrix size=256×256, slice thickness=0.6 mm, slice gap=0.1 mm, slice number=31.

#### Resting state fMRI data acquisition and analysis

Resting-state scans were collected using a gradient-echo echo-planner imaging (EPI) sequence developed in-house. This sequence consists of forward (top-down) and reverse (bottom-up) k-space trajectories. Geometric distortions in EPI images acquired using these two k-space trajectories feature distinct patterns, which were corrected using the approach described in (33). Scan parameters: TR=1500 ms, TE=15 ms, FOV=30×30 cm^2^, in-plane matrix size=80×80, slice thickness=0.6 mm, slice gap=0.1 mm, slice number=19.

fMRI data analyses procedures were previously reported (31). Geometric distortions in EPI images were first corrected using the data acquired using the forward and reverse k-space trajectory images (33). EPI time series were then subject to spatial independent component analysis (ICA) using the MELODIC package in FSL. Noise components that exhibit periodic temporal patterns resulting from respiration and cardiac pulsation were identified and removed in FSL. Images from one rat had the best positioning and were chosen as the template. EPI images were linearly co-registered to this template. The co-registered EPI images from all rats were subsequently averaged to generate a group template specific to this study and were subject to further co-registration to the group template. Time series were subject to second-order detrending along with voxel time-courses averaged from white matter and CSF as the nuance signal, low-pass filtered at 0.1 Hz. Voxel time courses from a mPFC seed region were averaged and served as the reference time course, which was cross correlated with time courses of the whole brain and subsequently subject to z transformation.

Functional connectivity data in TMS and sham hdTBS groups at two time points (early and late withdrawal) were subject to linear-mixed-effects modeling followed by post-hoc t-tests using the 3dLME function in AFNI. Multiple comparison corrections were carried out using the 3dCustSim function based on spatial autocorrelation function (ACF) fitted that to a model of the form(34):

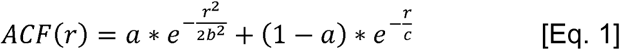

The outputs of the ACF function (parameters *a*, *b*, *c*) were fed into 3dClusSim to derive the cluster size for a given threshold (p<0.05, voxels in clusters>38). The above calculations were performed in AFNI (35). Note that Equation [1] does not assume a Gaussian distribution in spatial autocorrelation of MRI noise, leading to more robust estimate of the cluster size(34). Regions of interest (ROIs) within the activated clusters were mapped into anatomical regions based on a rat brain atlas (36). Voxel-wise functional connectivity values within individual ROIs were averaged and plotted.

## RESULTS

### hdTBS of mPFC prevented incubation of cocaine seeking

Previous studies have shown a critical role of mPFC in incubation of cocaine craving during abstinence (26). Thus, we tested whether TMS administration aimed at the mPFC during abstinence will prevent this incubation effect. Figure 1A illustrates the experimental timeline. A cocaine seeking test under extinction was conducted on abstinence day 1 such that lever-presses led to the delivery of a tone+light cue previously paired with cocaine infusions, but in the absence of cocaine. The rats were then divided into two groups: one received daily active hdTBS for 7 days (n=18) during abstinence day 14 to 20), while the other received sham hdTBS (n=15). A second cocaine seeking test was conducted on abstinence day 21 (11).

**Figure 1.**
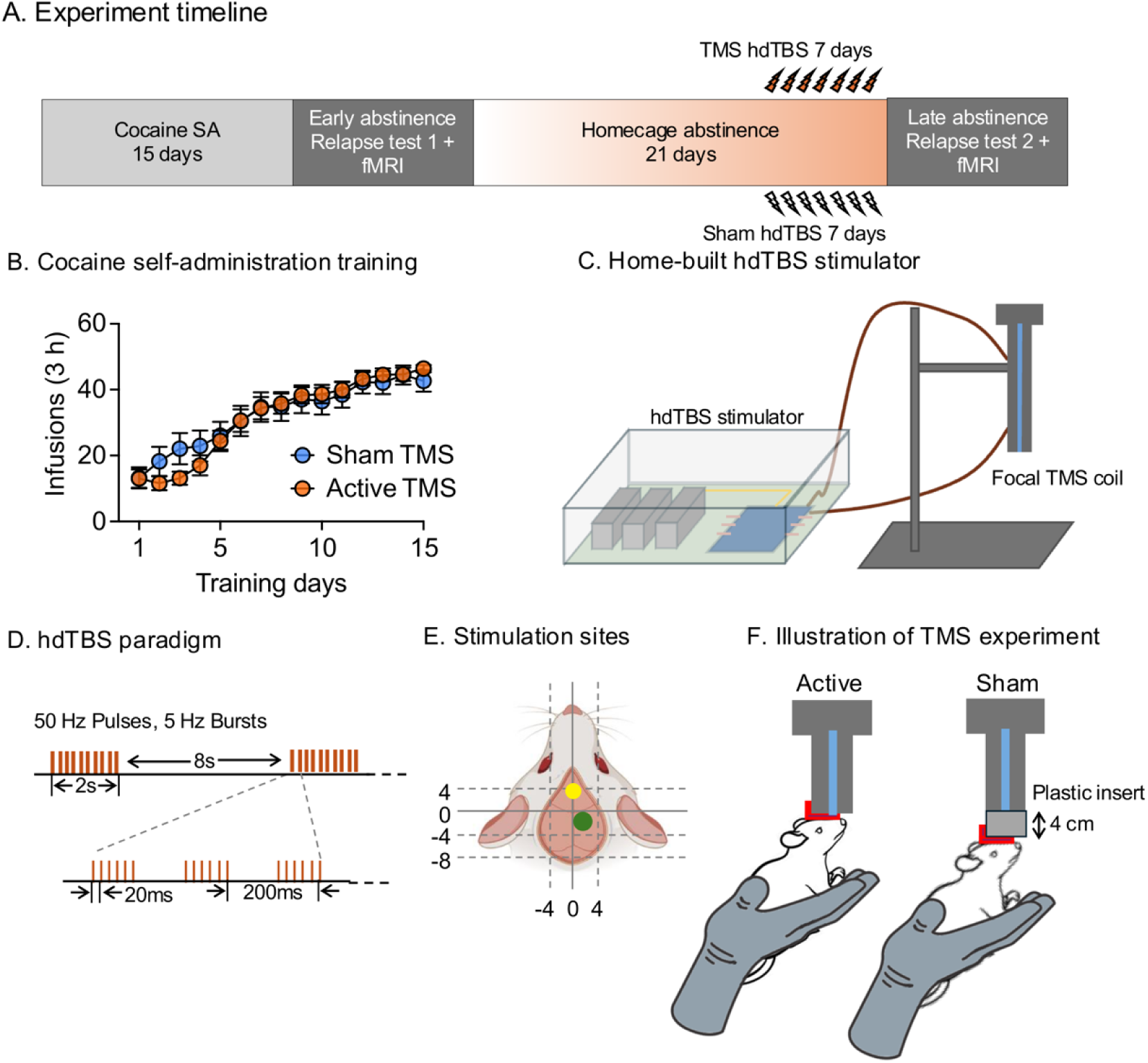
(**A**) *Experimental timeline:* Rats underwent cocaine self-administration on a fixed ratio 1 (FR-1) schedule for 15 days, followed by homecage abstinence. Cocaine seeking tests were conducted on abstinence days 1 and 21. Active or sham hdTBS was administered daily from day 14 to day 20. (**B**). Cocaine intake during self-administration training increased over time; there were no significant differences in cocaine infusions between the active TMS and sham groups (p = 0.67). (**C**) The coil was cooled using dry ice to prevent overheating. (**D**-**E**) Illustrations of the hdTBS procedure and TMS administration. The **green** and **yellow** areas indicate the hindlimb motor cortex and medial prefrontal cortex (mPFC), respectively. (**F**) For active TMS, rats were positioned with their heads against the coil surface and stimulated at 125% of motor threshold or maximum machine output if a rat’s motor threshold was too high. For sham TMS, the rat head was placed 4 cm below the coil and stimulated at 75% motor threshold.

**Figure 2.**
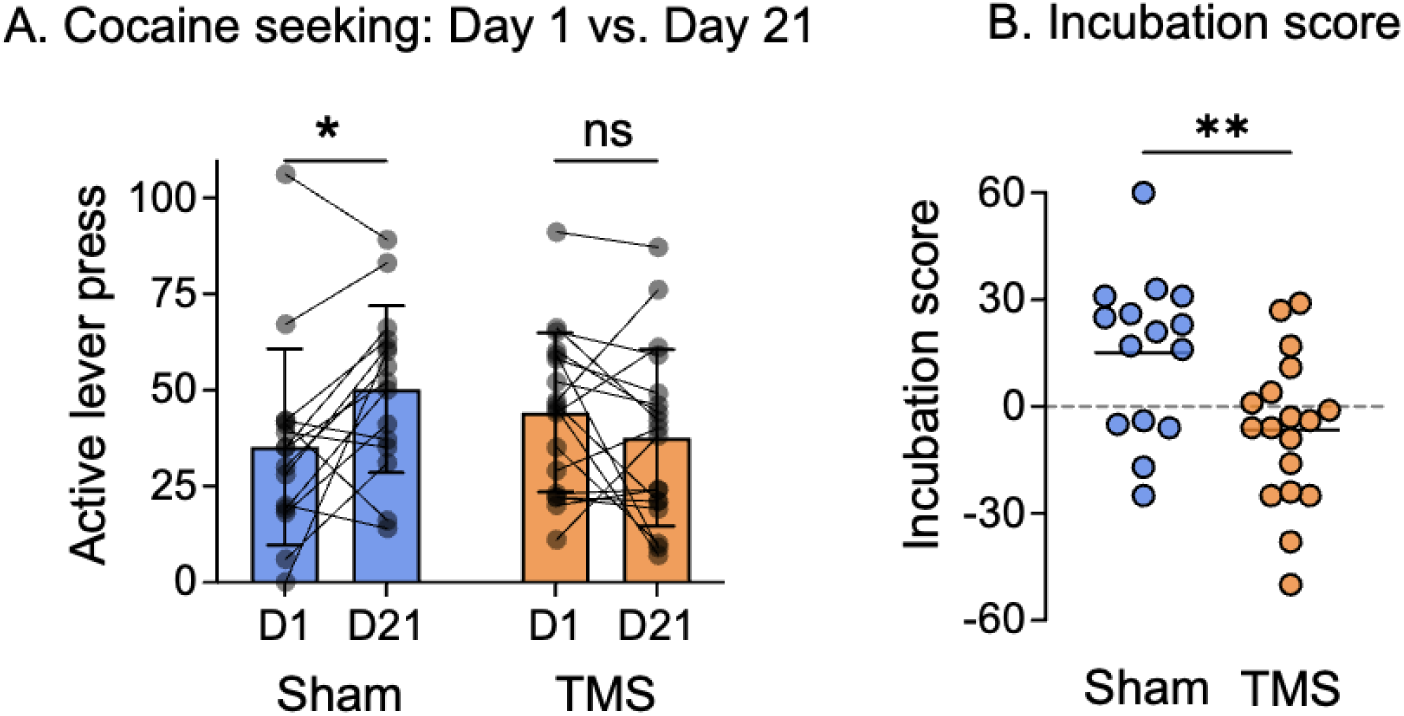
(**A**) Rats receiving sham TMS (n=15) showed a significant increase in cocaine seeking from day 1 (D1) to day 21 (D21), whereas no significant change was observed in rats treated with active hdTBS TMS (n=18). (**B**) Incubation score, defined as the difference in cocaine seeking between D21 and D1, was significantly higher in the sham TMS group than in the active hdTBS TMS group. *, *p=*0.047; **, *p* = 0.0069 (Tukey post-hoc test). There was no significant difference in D1 seeking (*p*=0.67).

LME modeling with training day and treatment group (active vs sham hdTBS) as fixed effects and individual rats as a random effect demonstrated that cocaine infusions increased across training days (F(14, 420)=51.27, *p*<0.0001); there were no significant group differences in cocaine SA prior to TMS treatment (F(1, 30)=0.039, *p*=0.84) Figure 1B. Figure 1C illustrates the rodent-specific focal TMS coil. Each hdTBS session delivers a total of 1200 pulses within the 200-s epoch. See Figure 1D for the hdTBS paradigm and 1E for TMS stimulation sites.

LME model analysis demonstrated a significant interaction between TREATMENT and TIME (F(_1,31_)=8.36, *p*=0.0069). Post-hoc analyses indicated a significant increase in cocaine seeking in the sham group on abstinence day 21 compared to day 1 (Figure 2A; Tukey-adjusted *p*=0.047), consistent with an incubation effect. In contrast, the active TMS group did not show a significant difference in cocaine-seeking behavior between abstinence day 21 and day 1. Furthermore, there was no significant difference in cocaine-seeking behavior between the active TMS and sham groups on abstinence day 1 (Tukey-adjusted, *p*=0.67), indicating comparable baseline levels. To further quantify changes in cocaine seeking over time, an incubation score was calculated for each subject as the difference in cocaine seeking between withdrawal day 21 and day 1. Results are shown in Figure 2B.

### hdTBS of mPFC reversed functional connectivity decreases during abstinence

Having established that a 7-day course of daily hdTBS prevented the incubation of cocaine craving, we next examine how brain function within the mPFC network changed during abstinence, and how these changes are influenced by hdTBS. Resting-state fMRI scans were conducted on the days following the first and second cocaine seeking tests (i.e., abstinence days 2 and 22). Due to technical reasons (see Methods), only a subset of rats (n=19) successfully completed both scans (n=10, active hdTBS; n=9, sham hdTBS).

Voxel-wise functional connectivity analyses (37) with seeds placed in the bilateral anterior prelimbic cortex (Supplemental Figure 3), demonstrated a significant Group x TIME interaction in three brain regions (right prelimbic cortex, right bed nucleus of the stria terminalis (BNST), and left ventrolateral thalamic nuclei (VL), *p*<0.05, corrected for multiple comparisons, Figure 3A). Functional connectivity plots for individual rats in these regions across both intervention groups and both time points are presented in Figure 3B and suggest that the interaction is driven by a decrease in FC in the sham but not active group across time.

**Figure 3.**
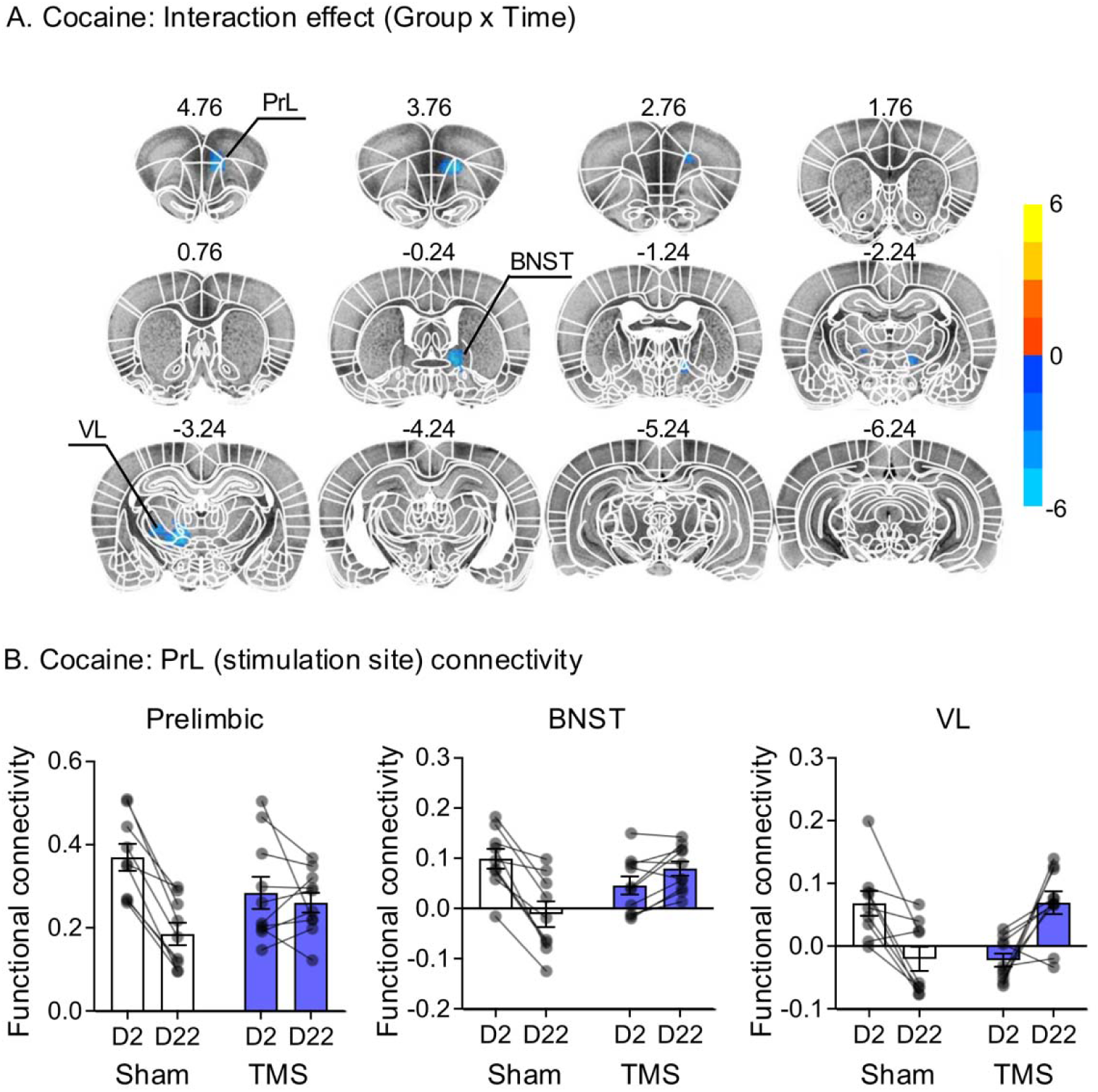
(**A**) Linear mixed-effects modeling showing significant interaction effect of Group x Time (TMS/Sham group x early/late abstinence) in 3 brain regions: PrL, thalamus and BNST. Group sizes: TMS (n=10), Sham (n=9). corrected p<0.05 (p<0.01 uncorrected). (**B**) Individual rat functional connectivity data for 3 regions of interest, using the prelimbic cortex (PrL) as the seed region (AP +3.76 and +4.76 mm) . Abbreviations: PrL, prelimbic cortex; BNST, bed nucleus of the stria terminalis; VL, ventrolateral nucleus of thalamus.

We next compared FC between early and late abstinence days within each TREATMENT condition, with the same prelimbic seed. In rats that received sham TMS, a significant reduction in FC was observed across multiple brain regions, including the orbital frontal cortex (OFC), cingulate cortex, prelimbic cortex (PrL), and nucleus accumbens (NAc) (Figure 4A). In contrast, rats treated with active hdTBS showed no significant functional connectivity reduction in these regions (Figure 4B). Functional connectivity plots from individual rats are shown in Figure 4C. These results demonstrate that abstinence from cocaine leads to a significant decrease in mPFC based circuits, which are reversed by hdTBS intervention during abstinence.

**Figure 4.**
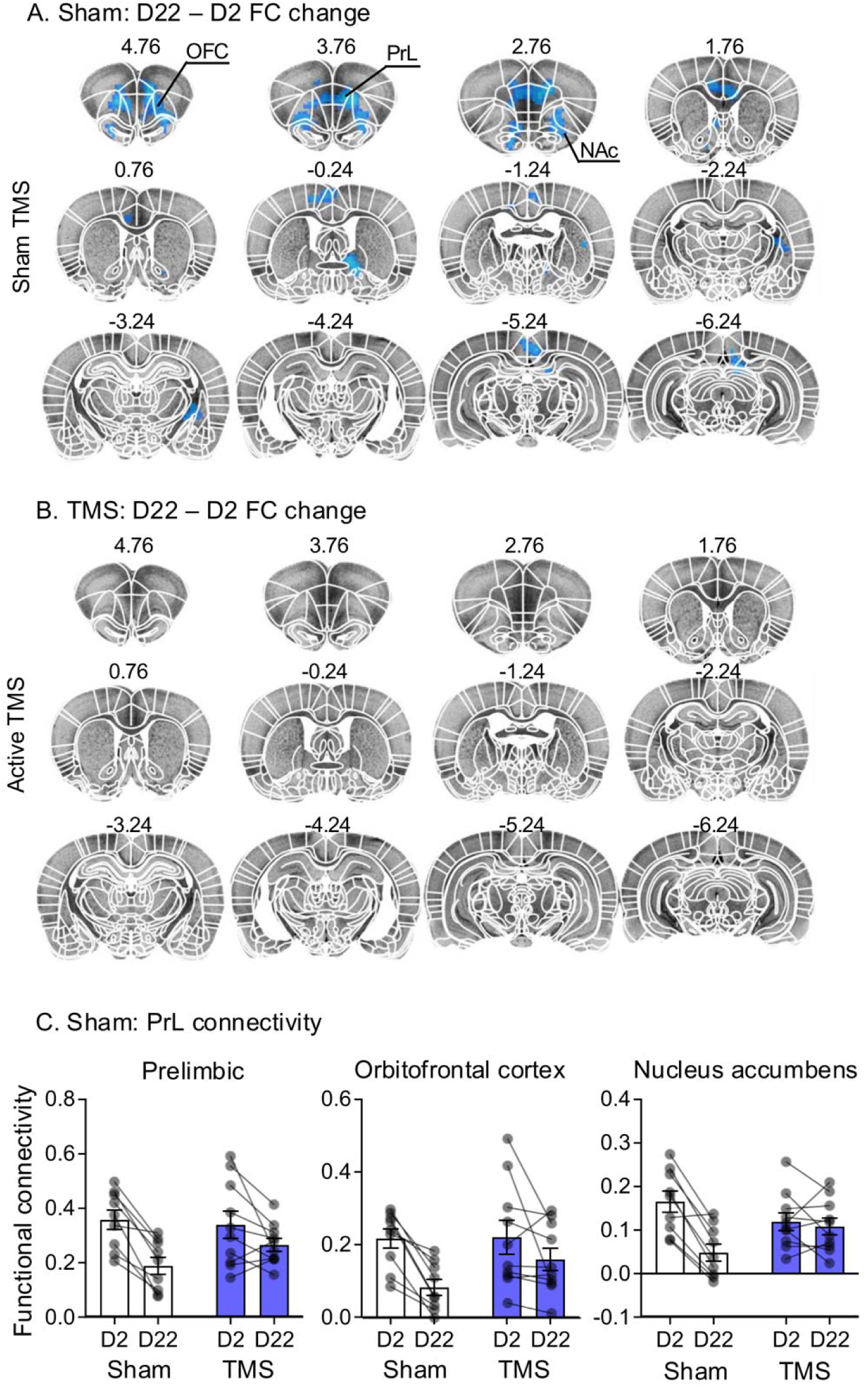
Functional connectivity during early (day 2, D2) vs. late (day 22, D22) abstinence. (**A**) Post-hoc t-test following linear mixed-effects modeling indicates a reduction in functional connectivity over time in the sham TMS group, particularly in the ACC, OFC, PrL, and NAc. (**B**) Seven days of daily session hdTBS reversed this decrease. (**C**) Individual rats’ function connectivity data for 3 regions of interest, using the prelimbic cortex (PrL) as the seed region. Group sizes: TMS (n=10), Sham (n=9). corrected p < 0.05 (p < 0.01 uncorrected). Abbreviations: OFC, orbitofrontal cortex; PrL, prelimbic cortex; NAc, nucleus accumbens; ACC, anterior cingulate cortex.

## DISCUSSION

We applied TMS using an hdTBS protocol to the mPFC of rats following cocaine self-administration. Daily hdTBS sessions over 7 days prevented the emergence of incubation of cocaine craving and restored functional connectivity between the PrL and OFC and NAc. The hdTBS protocol was well tolerated. Rats showed no changes in normal behaviors such as eating, drinking, or grooming, and no TMS-induced seizures were observed during or after stimulation. Behavioral performance deficits are unlikely to explain the results, as lever-pressing in the hdTBS group on day 21 was comparable to lever-pressing on day 1, prior to stimulation. In a companion study (38), focal hdTBS applied to the rat ACC similarly reduced incubation of opioid craving following electric barrier-induced voluntary abstinence, a procedure distinct from the classical homecage abstinence (39). Together, these findings suggest therapeutic potential for TMS across different classes of addictive drugs and abstinence procedures.

In sham-treated rats, several cortical and corticostriatal circuits, including OFC, ACC, PrL, and NAc, showed reduced functional connectivity with the PrL seed (Figures 3-4). These findings are consistent with prior human studies demonstrating decreased corticostriatal connectivity in SUD patients (40), as well as preclinical evidence implicating neuroadaptations in prefrontal-NAc glutamatergic pathways in relapse-related behavior (41,42). For example, Ma et al. (43) showed that synaptic neuroadaptations in the PrL projections to the NAc are critical to incubation of cocaine craving. More broadly, converging evidence indicates that prefrontal regions such as the PrL and OFC contribute to incubation and relapse-related behaviors (44–46). For instance, inactivation of the lateral OFC, the putative functional homolog of the human medial OFC, decreases both cue- and context-induced cocaine seeking (44,47,48). The connectivity reductions observed in our study may reflect impaired top-down regulation (49–51). The reversal of these connectivity reductions by hdTBS suggests a causal role for disrupted functional connectivity in incubation of cocaine seeking.

Additional reductions in connectivity in the sham group were observed between the PrL and BNST, as well as the PrL and VL thalamus. This pattern is notable because the BNST, a component of the extended amygdala, has been implicated in stress-induced reinstatement and negative affect during abstinence (52–54). Although the role of the VL thalamus in drug seeking is unknown, emerging evidence suggests that motor-related cortical systems like the secondary motor cortex contribute to incubation of cocaine craving (55,56).

Our study also incorporates several technical advances. First, the rat-specific TMS coil achieves a stimulation focality of approximately 2 mm (21,29), enabling targeted activation of specific brain systems with minimal off-target effects. This level of precision is comparable to that used clinically. Second, the hdTBS protocol enhances after-effects by 92% compared to conventional iTBS (21). Third, TMS was delivered in awake rats (29), eliminating confounds associated with anesthesia (57), an important consideration for repeated stimulation procedures.

Overall, our findings suggest that TMS inhibits relapse-related behaviors by restoring functional connectivity across distributed circuits involved in drug seeking and highlight potential targets for circuit-based neuromodulation. Given the well-established safety profile of TMS in humans and the cross-species applicability of fMRI, our findings have translational relevance for CUD, for which FDA-approved pharmacotherapies do not exist.

### Translating TMS therapy from rat models of cocaine dependence to CUD patients

Our TMS protocol targeted the mPFC, engaging both the PrL and anterior cingulate cortex. The rodent’s PrL is considered functionally analogous to the human dorsolateral prefrontal cortex (dlPFC) (51,58), a region central to executive control (59). Human imaging studies have reported hypofrontality, reduced prefrontal basal metabolism, in individuals with CUD patients (60,61). These deficits are thought to contribute to compulsive drug use despite negative consequences (49,50). Consistent with this idea, open-label TMS studies targeting the dlPFC have shown promising results in addiction treatment (3,6).

According to Vogt (62), the rodent cingulate cortex is homologous to Brodmann areas 24, 25, and 32 in humans, encompassing ventral anterior, subgenual, and dorsal anterior cingulate regions. Imaging studies have demonstrated ACC activation in response to drug-associated cues in individuals with CUD (63,64). A recent lesion network mapping study identified the paracingulate gyrus as a key hub associated with addiction remission(65). In line with these findings, our simultaneous engagement of ACC- and PrL-related circuits may have contributed to the observed reduction in incubation of cocaine seeking.

Together, based on (1) circuit-level models of addiction identifying the cingulate cortex as a central hub(65), (2) our findings that combined stimulation of ACC and PrL reduces relapse-like behavior, and (3) cross-species homologies between rodent and human prefrontal regions, we propose simultaneous targeting of the dlPFC and ACC (Brodmann areas 24, 25, and 32) as a neuromodulation strategy for CUD. Multi-site stimulation approaches are technically feasible (66) and warrant further investigation.

## Limitations

Our study has several limitations. First, only male rats were included to reduce variability associated with sex differences in brain anatomy and functional connectivity (67,68). In addition, our fMRI protocol requires light anesthesia (low-dose isoflurane and dexmedetomidine), and both sex- and estrous cycle-dependent responses to these agents have been reported (69,70), which would confound data interpretation. Hormonal fluctuations across the ovarian cycle are also known to influence functional connectivity in humans (71,72). Future studies should evaluate hdTBS effects in female rats.

Second, the durability of the TMS effect was not assessed. Stimulation was administered during abstinence days 14-20, and cocaine seeking was measured on day 21. Given that incubation of drug seeking persists for many months in this model (11,73), future studies should assess relapse to cocaine seeking at a longer time window after TMS treatment.

Third, while rodent models provide valuable insight, they do not fully capture the complexity of human addiction (74). Clinical relapse is triggered by multiple factors, including stress, drug cues, and drug re-exposure (9). Although the incubation model has translational relevance for cocaine, nicotine, and alcohol use (13–15,26), it primarily captures cue-induced craving and does not encompass other aspects of relapse. Future work should examine TMS effects on stress- and drug-induced relapse.

## Conclusions

In summary, focal hdTBS applied to the rat mPFC prevented incubation of cocaine craving and restored prefrontal functional connectivity. Given the established safety of TMS and its clinical applicability, our findings support its potential as a circuit-based intervention for CUD and related substance use disorders.

## Data and materials availability

All data used to evaluate the conclusions in the paper are present in the paper. Additional data related to this paper are available upon request from the corresponding author.

## Author’s contributions

SH, YD, ZM, AC, TS, OV and HL carried out the experiments; SH, YS, ZM, AC, TS, OV and HL performed data analysis. HL, EAS, ZX, YS and YY designed the study and HL, YD, YS and YY wrote the manuscript with feedback from co-authors. HN, MMH and HL developed the hdTBS stimulator hardware and software. All authors critically reviewed the content and approved the final version before submission.

## Acknowledgements

We gratefully acknowledge Dr. Khaled Moussawi during the early stages of this project for his insightful discussions and valuable suggestions on the administration and evaluation of TMS effects in rat models of cocaine addiction.

## References

1. United Nations Office on Drugs and Crime (2023): Global Report on Cocaine 2023: Local Dynamics, Global Challenges. UN.

2. Garnett MF, Miniño AM (2024): Drug Overdose Deaths in the United States, 2003–2023.

3. Ekhtiari H, Tavakoli H, Addolorato G, Baeken C, Bonci A, Campanella S, et al. (2019): Transcranial electrical and magnetic stimulation (tES and TMS) for addiction medicine: A consensus paper on the present state of the science and the road ahead. Neuroscience & Biobehavioral Reviews 104: 118–140.

4. Luigjes J, Segrave R, de Joode N, Figee M, Denys D (2019): Efficacy of Invasive and Non-Invasive Brain Modulation Interventions for Addiction. Neuropsychol Rev 29: 116–138.

5. Hanlon CA, Dowdle LT, Moss H, Canterberry M, George MS (2016): Mobilization of Medial and Lateral Frontal-Striatal Circuits in Cocaine Users and Controls: An Interleaved TMS/BOLD Functional Connectivity Study [no. 13], 2016/07/04 ed. Neuropsychopharmacology 41: 3032–3041.

6. Steele VR, Maxwell AM (2021): Treating cocaine and opioid use disorder with transcranial magnetic stimulation: A path forward. Pharmacology Biochemistry and Behavior 209: 173240.

7. Dinur-Klein L, Dannon P, Hadar A, Rosenberg O, Roth Y, Kotler M, Zangen A (2014): Smoking cessation induced by deep repetitive transcranial magnetic stimulation of the prefrontal and insular cortices: a prospective, randomized controlled trial. Biol Psychiatry 76: 742–749.

8. O’Brien CP (1997): Progress in the science of addiction [no. 9], 1997/09/01 ed. Am J Psychiatry 154: 1195–7.

9. O’Brien CP, Childress AR, McLellan AT, Ehrman R (1992): Classical conditioning in drug-dependent humans. Ann N Y Acad Sci 654: 400–415.

10. Grimm JW, Hope BT, Wise RA, Shaham Y (2001): Neuroadaptation - Incubation of cocaine craving after withdrawal [no. 6843]. Nature 412: 141–142.

11. Lu L, Grimm JW, Hope BT, Shaham Y (2004): Incubation of cocaine craving after withdrawal: a review of preclinical data. Neuropharmacology 47 Suppl 1: 214–26.

12. Neisewander JL, Baker DA, Fuchs RA, Tran-Nguyen LT, Palmer A, Marshall JF (2000): Fos protein expression and cocaine-seeking behavior in rats after exposure to a cocaine self-administration environment [no. 2]. J Neurosci 20: 798–805.

13. Bedi G, Preston KL, Epstein DH, Heishman SJ, Marrone GF, Shaham Y, de Wit H (2011): Incubation of cue-induced cigarette craving during abstinence in human smokers [no. 7]. Biol Psychiatry 69: 708–11.

14. Li P, Wu P, Xin X, Fan YL, Wang GB, Wang F, et al. (2015): Incubation of alcohol craving during abstinence in patients with alcohol dependence [no. 3]. Addict Biol 20: 513–22. doi: 10.1111/adb.12140. Epub 2014 Apr 2.

15. Parvaz MA, Moeller SJ, Goldstein RZ (2016): Incubation of Cue-Induced Craving in Adults Addicted to Cocaine Measured by Electroencephalography [no. 11]. JAMA Psychiatry 73: 1127–1134. doi: 10.1001/jamapsychiatry.2016.2181.

16. Deng Z-D, Lisanby SH, Peterchev AV (2013): Electric field depth–focality tradeoff in transcranial magnetic stimulation: simulation comparison of 50 coil designs. Brain Stimul 6: 1–13.

17. Cohen SL, Bikson M, Badran BW, George MS (2022): A visual and narrative timeline of US FDA milestones for Transcranial Magnetic Stimulation (TMS) devices [no. 1]. Brain Stimul 15: 73–75.

18. Wilson MT, Tang AD, Iyer K, McKee H, Waas J, Rodger J (2018): The challenges of producing effective small coils for transcranial magnetic stimulation of mice [no. 3]. Biomed Phys Eng Express 4: 037002.

19. Meng Q, Jing L, Badjo JP, Du X, Hong E, Yang Y, et al. (2018): A novel transcranial magnetic stimulator for focal stimulation of rodent brain. Brain Stimul 11: 663–665.

20. Huang YZ, Edwards MJ, Rounis E, Bhatia KP, Rothwell JC (2005): Theta burst stimulation of the human motor cortex [no. 2]. Neuron 45: 201–6.

21. Meng Q, Nguyen H, Vrana A, Baldwin S, Li CQ, Giles A, et al. (2022): A high-density theta burst paradigm enhances the aftereffects of transcranial magnetic stimulation: Evidence from focal stimulation of rat motor cortex. Brain Stimul 15: 833–842.

22. Moorman DE, Aston-Jones G (2023): Prelimbic and infralimbic medial prefrontal cortex neuron activity signals cocaine seeking variables across multiple timescales. Psychopharmacology 240: 575–594.

23. Fuchs RA, Evans KA, Ledford CC, Parker MP, Case JM, Mehta RH, See RE (2005): The role of the dorsomedial prefrontal cortex, basolateral amygdala, and dorsal hippocampus in contextual reinstatement of cocaine seeking in rats [no. 2]. Neuropsychopharmacology 30: 296–309.

24. West EA, Saddoris MP, Kerfoot EC, Carelli RM (2014): Prelimbic and infralimbic cortical regions differentially encode cocaine-associated stimuli and cocaine-seeking before and following abstinence. European Journal of Neuroscience 39: 1891–1902.

25. McFarland K, Lapish CC, Kalivas PW (2003): Prefrontal glutamate release into the core of the nucleus accumbens mediates cocaine-induced reinstatement of drug-seeking behavior. [no. 8]. J Neurosci 23: 3531–7.

26. Chow JJ, Pitts KM, Negishi K, Madangopal R, Dong Y, Wolf ME, Shaham Y (2025): Neurobiology of the incubation of drug craving: An update. Pharmacol Rev 77: 100022.

27. Ma Z, Duan Y, Fredriksson I, Tsai P-J, Batista A, Lu H, et al. (2024): Role of dorsal striatum circuits in relapse to opioid seeking after voluntary abstinence. Neuropsychopharmacol 1–9.

28. Bates D, Mächler M, Bolker B, Walker S (2015): Fitting Linear Mixed-Effects Models Using lme4. Journal of Statistical Software 67: 1–48.

29. Cermak S, Meng Q, Peng K, Baldwin S, Mejías-Aponte CA, Yang Y, Lu H (2020): Focal transcranial magnetic stimulation in awake rats: enhanced glucose uptake in deep cortical layers. J Neurosci Methods 339: 108709.

30. Brynildsen JK, Hsu LM, Ross TJ, Stein EA, Yang Y, Lu H (2016): Physiological characterization of a robust survival rodent fMRI method. Magn Reson Imaging. 10.1016/j.mri.2016.08.010

31. Lu H, Zou Q, Gu H, Raichle ME, Stein EA, Yang Y (2012): Rat brains also have a default mode network. Proceedings of the National Academy of Sciences 109: 3979–3984.

32. Lu H, Xi ZX, Gitajn L, Rea W, Yang Y, Stein EA (2007): Cocaine-induced brain activation detected by dynamic manganese-enhanced magnetic resonance imaging (MEMRI). [no. 7]. Proc Natl Acad Sci U S A 104: 2489–94.

33. Chang H, Fitzpatrick J (1992): A technique for accurate magnetic-resonance-imaging in the presence of field inhomogeneities [no. 3]. Ieee Transactions on Medical Imaging 11: 319–329.

34. Cox RW, Chen G, Glen DR, Reynolds RC, Taylor PA (2017): FMRI Clustering in AFNI: False-Positive Rates Redux. Brain Connectivity 7: 152–171.

35. Cox R (1996): AFNI: software for analysis and visualization of functional magnetic resonance neuroimages. [no. 3]. Comput Biomed Res 29: 162–73.

36. Paxinos G, Watson C (2007): The Rat Brain in Stereotaxic Coordinates, 6th ed. Elsevier Inc.

37. Biswal B, Yetkin FZ, Haughton VM, Hyde JS (1995): Functional connectivity in the motor cortex of resting human brain using echo-planar MRI. [no. 4]. Magn Reson Med 34: 537–41.

38. Ma Z, Duan Y, Nguyen H, Lin S, Haque MM, Wang D, et al. (2026, March 6): Focal Transcranial Magnetic Stimulation of the Rat Anterior Cingulate Cortex Inhibits Incubation of Opioid Craving after Voluntary Abstinence. bioRxiv, p 2026.03.04.709400.

39. Fredriksson I, Venniro M, Reiner DJ, Chow JJ, Bossert JM, Shaham Y (2021): Animal Models of Drug Relapse and Craving after Voluntary Abstinence: A Review. Pharmacol Rev 73: 1050–1083.

40. Motzkin JC, Baskin-Sommers A, Newman JP, Kiehl KA, Koenigs M (2014): Neural correlates of substance abuse: Reduced functional connectivity between areas underlying reward and cognitive control. Hum Brain Mapp 35: 4282–4292.

41. Marchant NJ, Kaganovsky K, Shaham Y, Bossert JM (2015): Role of corticostriatal circuits in context-induced reinstatement of drug seeking. Brain Res 1628: 219–232.

42. McGlinchey EM, James MH, Mahler SV, Pantazis C, Aston-Jones G (2016): Prelimbic to Accumbens Core Pathway Is Recruited in a Dopamine-Dependent Manner to Drive Cued Reinstatement of Cocaine Seeking. J Neurosci 36: 8700–8711.

43. Ma Y-Y, Lee BR, Wang X, Guo C, Liu L, Cui R, et al. (2014): Bidirectional modulation of incubation of cocaine craving by silent synapse-based remodeling of prefrontal cortex to accumbens projections. Neuron 83: 1453–1467.

44. Lasseter HC, Ramirez DR, Xie X, Fuchs RA (2009): Involvement of the Lateral Orbitofrontal Cortex in Drug Context-induced Reinstatement of Cocaine-seeking Behavior in Rats. Eur J Neurosci 30: 1370–1381.

45. Mesa JR, Wesson DW, Schwendt M, Knackstedt LA (2022): The roles of rat medial prefrontal and orbitofrontal cortices in relapse to cocaine-seeking: A comparison across methods for identifying neurocircuits. Addict Neurosci 4: 100031.

46. Fanous S, Goldart EM, Theberge FRM, Bossert JM, Shaham Y, Hope BT (2012): Role of orbitofrontal cortex neuronal ensembles in the expression of incubation of heroin craving. J Neurosci 32: 11600–11609.

47. Fuchs R, Evans K, Parker M, See R (2004): Differential involvement of orbitofrontal cortex subregions in conditioned cue-induced and cocaine-primed reinstatement of cocaine seeking in rats. [no. 29]. J Neurosci 24: 6600–10.

48. Lasseter HC, Xie X, Ramirez DR, Fuchs RA (2010): Prefrontal cortical regulation of drug seeking in animal models of drug relapse. Curr Top Behav Neurosci 3: 101–117.

49. Goldstein RZ, Volkow ND (2011): Dysfunction of the prefrontal cortex in addiction: neuroimaging findings and clinical implications [no. 11]. Nat Rev Neurosci 12: 652–669.

50. Jentsch JD, Taylor JR (1999): Impulsivity resulting from frontostriatal dysfunction in drug abuse: implications for the control of behavior by reward-related stimuli. [no. 4]. Psychopharmacology (Berl) 146: 373–90.

51. Koob GF, Volkow ND (2016): Neurobiology of addiction: a neurocircuitry analysis [no. 8]. Lancet Psychiatry 3: 760–73.

52. Smith RJ, Aston-Jones G (2008): Noradrenergic transmission in the extended amygdala: role in increased drug-seeking and relapse during protracted drug abstinence. Brain Struct Funct 213: 43–61.

53. Leri F, Flores J, Rodaros D, Stewart J (2002): Blockade of stress-induced but not cocaine-induced reinstatement by infusion of noradrenergic antagonists into the bed nucleus of the stria terminalis or the central nucleus of the amygdala. J Neurosci 22: 5713–5718.

54. Shalev U, Highfield D, Yap J, Shaham Y (2000): Stress and relapse to drug seeking in rats: studies on the generality of the effect. Psychopharmacology (Berl) 150: 337–346.

55. Gómez-Pineda V, Huang D, Chen Y, Guo C, Ma Y-Y (2026): Contrasting roles of secondary motor cortex projections to the dorsolateral and dorsomedial striatum in incubation of cocaine-seeking. Neuropsychopharmacology. 10.1038/s41386-026-02385-3

56. Huang D, Ma Y-Y (2025): Downregulation of Small-Conductance Potassium Channels in Cortical Pyramidal Neurons of the Supplementary Motor Cortex Underlies High Cocaine-Seeking Behaviors. Biol Psychiatry S0006-3223(25)01607–5.

57. Gersner R, Kravetz E, Feil J, Pell G, Zangen A (2011): Long-term effects of repetitive transcranial magnetic stimulation on markers for neuroplasticity: differential outcomes in anesthetized and awake animals [no. 20]. J Neurosci 31: 7521–6.

58. Jasinska AJ, Chen BT, Bonci A, Stein EA (2015): Dorsal medial prefrontal cortex (MPFC) circuitry in rodent models of cocaine use: implications for drug addiction therapies. Addiction Biology 20: 215–226.

59. Goldstein RZ, Volkow ND (2002): Drug addiction and its underlying neurobiological basis: neuroimaging evidence for the involvement of the frontal cortex [no. 10], 2002/10/03 ed. Am J Psychiatry 159: 1642–52.

60. Volkow N, Mullani N, Gould K, Adler S, Krajewski K (1988): Cerebral blood-flow in chronic cocaine users - a study with positron mission tomography. British Journal of Psychiatry 641–648.

61. Volkow ND, Hitzemann R, Wang G-J, Fowler JS, Wolf AP, Dewey SL, Handlesman L (1992): Long-Term frontal brain metabolic changes in cocaine abusers. Synapse 11: 184–190.

62. Vogt BA (n.d.): Cingulate cortex and pain architecture. The Rat Nervous System, Fourth. Elsevier Inc., pp 575–578.

63. Garavan H, Pankiewicz J, Bloom A, Cho JK, Sperry L, Ross TJ, et al. (2000): Cue-induced cocaine craving: neuroanatomical specificity for drug users and drug stimuli. [no. 11]. Am J Psychiatry 157: 1789–98.

64. Childress AR, Mozley PD, McElgin W, Fitzgerald J, Reivich M, O’Brien CP (1999): Limbic activation during cue-induced cocaine craving [no. 1], 1999/01/19 ed. Am J Psychiatry 156: 11–8.

65. Joutsa J, Moussawi K, Siddiqi SH, Abdolahi A, Drew W, Cohen AL, et al. (2022): Brain lesions disrupting addiction map to a common human brain circuit. Nat Med 28: 1249–1255.

66. Sinisalo H, Rissanen I, Kahilakoski O-P, Souza VH, Tommila T, Laine M, et al. (2024): Modulating brain networks in space and time: Multi-locus transcranial magnetic stimulation. Clinical Neurophysiology 158: 218–224.

67. Wang Z, Guo Y, Mayer EA, Holschneider DP (2019): Sex differences in insular functional connectivity in response to noxious visceral stimulation in rats. Brain Research 1717: 15–26.

68. Sumiyoshi A, Nonaka H, Kawashima R (2017): Sexual differentiation of the adolescent rat brain: A longitudinal voxel-based morphometry study. Neuroscience Letters 642: 168–173.

69. Vincent KF, Mallari OG, Dillon EJ, Stewart VG, Cho AJ, Dong Y, et al. (2023): Oestrous cycle affects emergence from anaesthesia with dexmedetomidine, but not propofol, isoflurane, or sevoflurane, in female rats. British Journal of Anaesthesia 131: 67–78.

70. Wasilczuk AZ, Rinehart C, Aggarwal A, Stone ME, Mashour GA, Avidan MS, et al. (2024): Hormonal basis of sex differences in anesthetic sensitivity. Proceedings of the National Academy of Sciences 121: e2312913120.

71. Filippi M, Valsasina P, Misci P, Falini A, Comi G, Rocca MA (2013): The organization of intrinsic brain activity differs between genders: A resting-state fMRI study in a large cohort of young healthy subjects. Human Brain Mapping 34: 1330–1343.

72. Pritschet L, Santander T, Taylor CM, Layher E, Yu S, Miller MB, et al. (2020): Functional reorganization of brain networks across the human menstrual cycle. NeuroImage 220: 117091.

73. Madangopal R, Drake OR, Pham DQ, Lennon VA, Weber SJ, Lee J, et al. (2026): Persistent vulnerability to heroin relapse across the adult lifespan in rats. bioRxiv 2026.03.18.712140.

74. Venniro M, Banks ML, Heilig M, Epstein DH, Shaham Y (2020): Improving translation of animal models of addiction and relapse by reverse translation. Nat Rev Neurosci 21: 625–643.

