## Supplemental Figure for "Targeted medial prefrontal cortex stimulation prevents incubation of cocaine craving and restores functional connectivity"

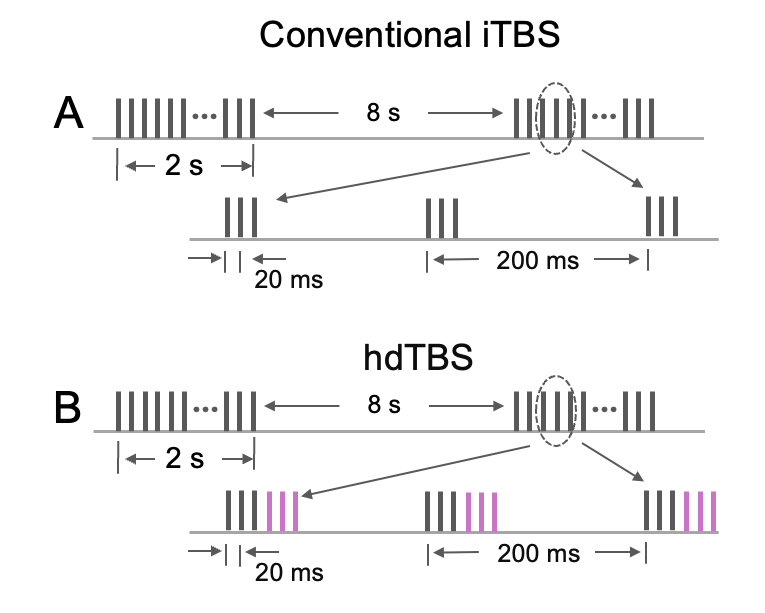

Figure S1. (A): The FDA-approved conventional intermittent theta burst stimulation (iTBS). Bursts of pulses are applied every 200 ms during the ON period; each consists of 3 pulses with an inter-pulse interval (IPI) of 20 ms (50 Hz), 600 pulses/session. (B): 50 Hz hdTBS, identical to (A) except that each burst consists of up to 6 pulses/burst, 1200 pulses/session.

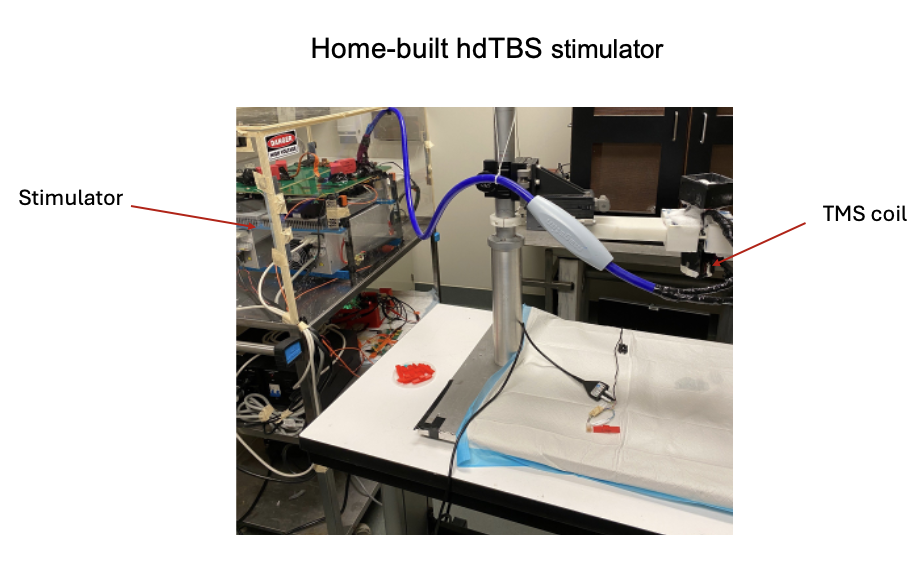

Figure S2. The hdTBS stimulator developed in house. The rat-specific TMS coil features vertical, asymmetric windings rather than in the conventional horizontal configuration and incorporates a small magnetic core to improve coil efficiency.

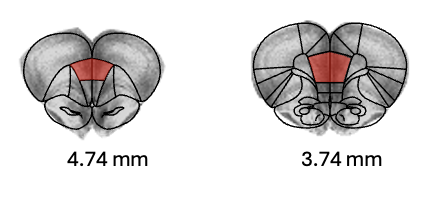

Figure S3. Illustration of seed regions used for resting state fMRI data analysis. The numbers below figures indicate coordinates relative to bregma.
